# Role of the medial agranular cortex in unilateral spatial neglect

**DOI:** 10.1101/2023.11.24.568203

**Authors:** Daisuke Ishii, Hironobu Osaki, Arito Yozu, Tatsuya Yamamoto, Satoshi Yamamoto, Mariko Miyata, Yutaka Kohno

**Author notes:** Corresponding author: Daisuke Ishii. These authors contributed equally: Daisuke Ishii and Hironobu Osaki.

## Abstract

Unilateral spatial neglect (USN) results from impaired attentional networks and can affect various sensory modalities, such as visual and somatosensory. The rodent medial agranular cortex (AGm), located in the medial part of the forebrain from rostral to caudal direction, is considered a region associated with spatial attention. The AGm selectively receives multisensory input with the rostral AGm receiving somatosensory input and caudal part receiving visual input. Our previous study showed slower recovery from neglect with anterior AGm lesion using the somatosensory neglect assessment. Conversely, the functional differences in spatial attention across the entire AGm locations (anterior, intermediate, and posterior parts) are unknown. Here, we investigated the relationship between the severity of neglect and various locations across the entire AGm in a mouse stroke model using a newly developed program-based analysis method that does not require human intervention. Among the various lesion positions, acute severity was higher with the lesion in the intermediate rostrocaudal position. On the other hand, the recovery from USN-like behavior after this phase tended to be slower in cases with more rostral lesions in the AGm. Additionally, no motor paralysis was observed in any of the mice with lesions in each AGm. These results suggest that the intermediate rostrocaudal position of the AGm may significantly influence selection of the direction, regardless of the areas to which it is connected. On the contrary, recovery from USN-like behavior may be dependent on the areas to which it is connected.

**Highlights:**

Lesion of the rodent medial agranular cortex (AGm) results in unilateral spatial neglect (USN).
In the acute phase, the severity was higher with lesions in the intermediate AGm position.
Recovery from somatosensory USN tended to be slower with rostral AGm lesions.
Recovery from USN may depend on sensory modalities associated with the connected areas.
Our results revealed location-dependent differences in attentional functions within the AGm.

## 1. Introduction

Unilateral spatial neglect (USN) is a symptom typically caused by damage to one side of the brain (either the right or left hemisphere) and characterized by ignoring or failing to perceive stimuli on the opposite side of the damaged brain hemisphere [1, 2]. Patients with USN have difficulty dressing, moving, and performing other activities of daily living and often require longer hospital stays than patients without neglect [3]. The development of animal models is important for developing effective rehabilitation strategies.

The rodent medial agranular cortex (AGm), located in the medial part of the forebrain from rostral to caudal direction, is considered a region that is associated with spatial attention [4–6]. The AGm has been reported to integrate various sensory information, including somatosensory, visual, and auditory information [7–9], together with motor actions and decision-making [10–12]. In a role of AGm in spatial attention function, our and previous studies have reported that damage to the AGm results in USN-like symptoms in visual, auditory, and somatosensory modalities [6, 13, 14].

A structural feature of the AGm is that the rostral AGm receives more somatosensory inputs, while the central and caudal AGm receives more inputs from the visual field [15, 16]. Our previous study revealed that within some AGm regions (around 1.5 mm from the bregma), recovery from neglect was slower in individuals with lesions located more anteriorly, as assessed by a somatosensory-dependent assessment [13]. However, the relationship between the sites of lesion across a broad range of the AGm (the intermediate, rostrocaudal, and posterior positions) and severity of neglect has not been investigated.

Therefore, we combined the photothrombotic stroke mouse model with automated USN-like behavior video analysis to extensively investigate the relationship between the location of AGm lesions and the severity of acute symptoms depending on somatosensory information. Additionally, we examined the differences in functional recovery from USN-like behavior with respect to AGm lesion location. Furthermore, motor deficits following AGm lesion have not been thoroughly examined and require comprehensive investigation. Moreover, we investigated the role of the AGm in motor function in detail using the ladder rung walking task.

## 2. Materials and Methods

### 2.1. Animals

All 9-week-old male C57BL/6J mice were purchased from Japan SLC Co., Shizuoka, Japan, and housed individually in cages under controlled conditions (12-h light/dark cycle, 23°C ± 1°C) with food and water *ad libitum*. For the evaluation of ipsilesional spatial bias resulting from cerebral infarction, we divided the animals into five groups: control (n = 14), AGm2.0 infarction (n = 6), AGm1.5 infarction (n = 18), AGm1.0 infarction (n = 14), and AGm0.5 infarction (n = 15) groups. Additionally, for the evaluation of motor paralysis resulting from cerebral infarction, we divided the animals into five groups: control (n = 11), AGm2.0 infarction (n = 15), AGm1.5 infarction (n = 10), AGm1.0 infarction (n = 8), and AGm0.5 infarction (n = 9) groups. All experiments were performed in accordance with the “Guidelines for the Proper Conduct of Animal Experiments” established by the Science Council of Japan in 2006, and were approved by the Animal Care and Use Committee of Ibaraki Prefectural University of Health Sciences (approval number 2022-14). This study is reported in accordance with ARRIVE guidelines.

### 2.2. Photothrombotic infarction in the AGm

In this study, we used focal photothrombotic infarction in the AGm as a stroke model [13, 17–19]. The detailed descriptions can be found in the previous study [13]. We positioned an optic fiber (diameter, 0.4 mm; 11 mW; Thorlabs Inc., Newton, NJ, USA) at the right AGm2.0, 1.5, 1.0, and 0.5 (2.0 mm, 1.5 mm, 1.0 mm, and 0.5 mm anterior to the bregma, respectively; all 0.6 mm lateral to the midline) (Figs. 1A and 1B). In the control group, mice received Rose Bengal by intraperitoneal injection. Subsequently, their scalps were sutured and they were returned to their home cages.

**Figure 1.**
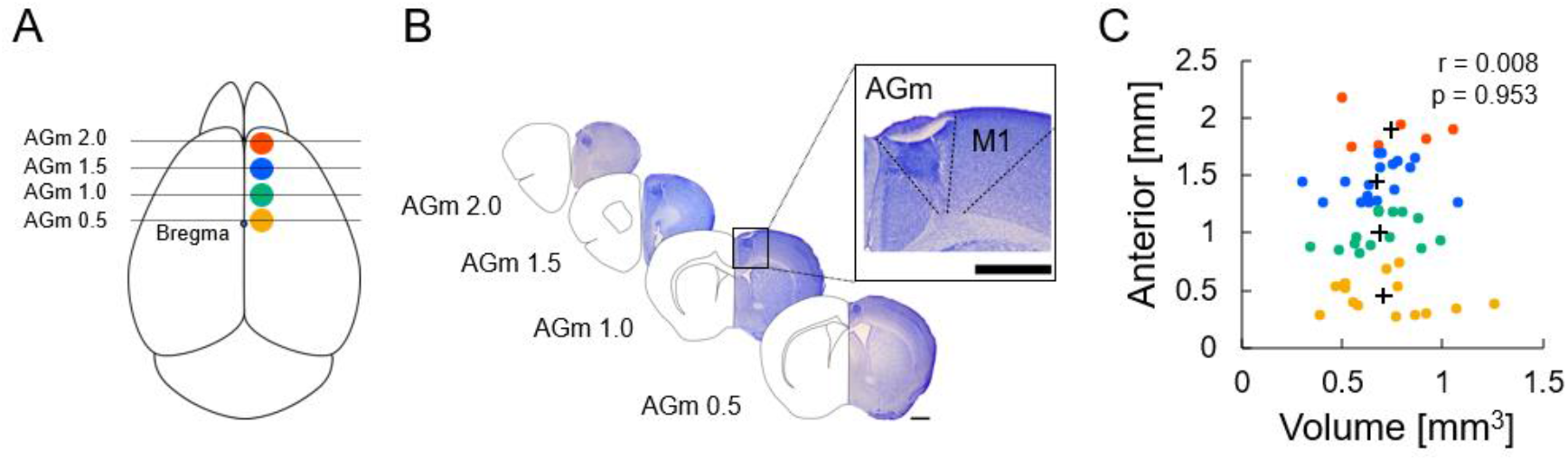
Photothrombotic infarction and evaluation of ipsilesional spatial bias. (A) Photothrombotic infarction was induced by local irradiation with a green light (2.0, 1.5, 1.0, and 0.4 mm anterior to bregma and 0.6 mm lateral to the midline). (B) The infarct site was identified by Nissl staining. Typical stions from the AGm2.0 (n = 6), AGm1.5 (n = 18), AGm1.0 (n = 14), and AGm0.5 groups (n = 15) are shown. The magnified typical lesion area and the M1 and AGm regions are represented by dotted lines. The dotted area in the AGm indicates the lesion site. Sections = 100 μm. Scale bar = 1 mm. (C) A scatterplot of the infarct site and volumes at postoperative day (POD) 18. + indicates the average centroid of the lesion area (A–P direction) in the AGm2.0, AGm1.5, AGm1.0, and AGm0.5 groups and average infarct volume in each group. Correlations were identified using the Pearson’s correlation coefficient. For nonlongitudinal analysis of the infarction volume, the statistical analysis of mean values was conducted using one-way analysis of variance (ANOVA).

### 2.3. Behavioral experiments

#### 2.3.1. Evaluation of ipsilesional spatial bias

To evaluate the ipsilesional spatial bias following a focal cerebral infarction, we used the eight-arm radial maze (Muromachi Kikai Co. Ltd, Tokyo, Japan) with a previously described protocol [13]. This assessment was performed in one prephotothrombosis test (pretest) and nine postphotothrombosis tests (Posttest). Two days before the pretest, mice were habituated by exposure to the maze in a sound-attenuated and dark room for 10 min. All mice underwent the pretest prior to photothrombosis. Two days thereafter, each mouse underwent a total of nine posttests. During the entire process, each mouse was placed individually in the center of an octagonal platform and was permitted to freely explore the maze for 11 min. We analyzed their behavior from 10 s after the mouse was placed in the eight-maze until 10 min and 10 s. Animal movements were recorded by an infrared digital video camera (the infrared filter was removed, C922; Logitech, Lausanne, Switzerland). At the end of the behavioral test, we removed the mouse and thoroughly cleaned the apparatus with water.

We calculated the left-side selection rate and severity of spatial bias (the left-side selection rate), and degree of recovery at POD4–7 and POD10–18 using a previously described protocol [13] (Fig.2A).

**Figure 2.**
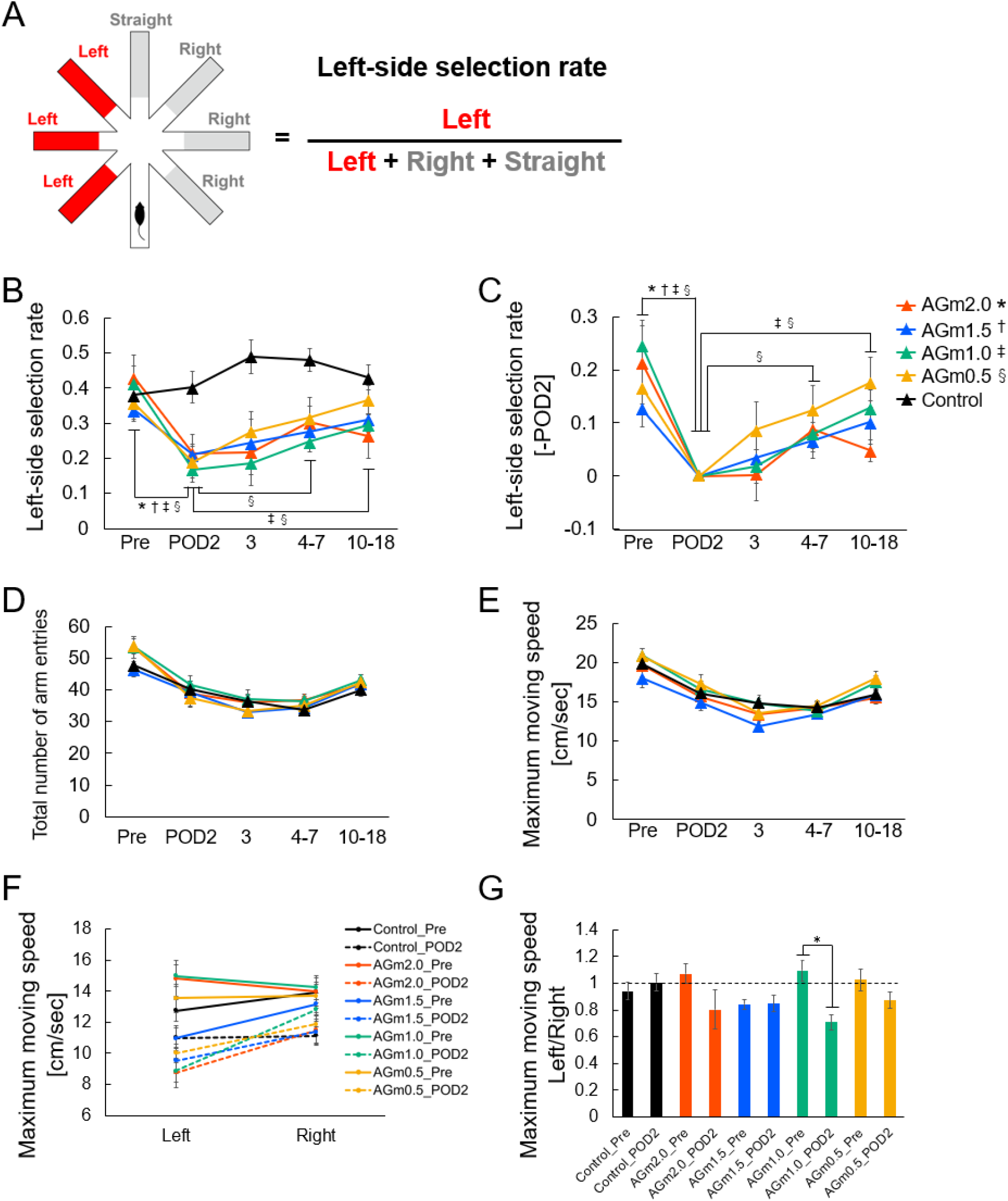
Changes to the left-side selection rate, total number of arm entries, and maximum moving speed after infarction. (A) An eight-arm radial maze was used. If the mouse was leaving the arm, the selection of Left Arm (Red) on the left-side of the maze with respect to the mouse was counted as a left-side selection, whereas the selection of Right Arm (gray) on the right-side of the maze in relation to the mouse was counted as a right-side selection. Selecting the Straight Arm (gray) in front of the mouse through the midline of the maze was counted as a straight selection. (B) Left-side selection rate during the pretest, POD2, POD3, POD4–7, and POD10–18 periods for the control group (n = 14), the AGm2.0 infarction group (n = 6), the AGm1.5 infarction group (n = 18), the AGm1.0 infarction group (n = 14) and the AGm0.5 infarction group (n = 15). *p < 0.05, ^†^p < 0.05, ^‡^p < 0.05, ^§^p < 0.05, POD2 vs. other time points in AGm2.0, 1.5, 1.0, and 0.5 groups, respectively according to two-way repeated measures ANOVA followed by Bonferroni’s test. (C) Changes in left-side selection rate with respect to POD2 for the AGm2.0, 1.5, 1.0, and 0.5 groups. (D) Total number of arm entries during the pretest, POD2, POD3, POD4–7, and POD10–18 periods. (E) Maximum moving speed among all arms during the pretest, POD2, POD3, POD4–7, and POD10–18 periods. (F) Maximum moving speed for the right and left selections at the pre and POD2 periods, and (G) left-right ratio of maximum moving speed for right and left selections at the pretest and POD2 periods. *p < 0.05, Pre vs. POD2 in AGm1.0 group by paired t-test. All data points indicate the mean ± SEM.

The left-side selection rate was determined by the following equation:

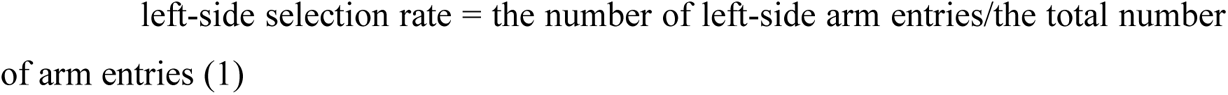

We defined severity as the magnitude of the decrease in left-sided selectivity rate on the first posttest (postoperative day 2; POD2) compared with the pretest, which was determined using the following formula:

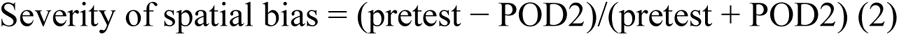

We defined the degree of recovery as the magnitude of the increase in the left-side selection rate during the average of POD4–7 or POD10–18 compared with POD2, and determined it using the following equations:

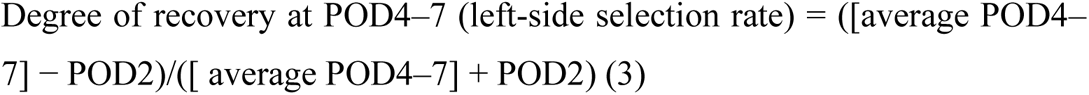

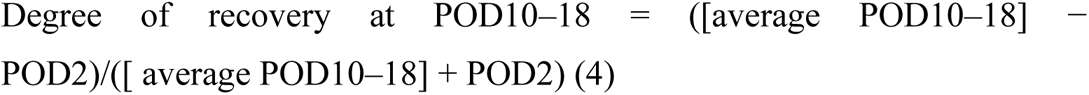

Additionally, we calculated the moving averages of severity, recovery and coordinates of the lesion sites using the movmean function in Matlab (R2021b, MathWorks, Natick, MA, United States). We determined the window size as the point where residuals decreased following an increase (the window size for assessing severity at POD2 was 10, and for recovery at POD4–7 and POD10–18 was 15, as shown in Supplementary Figs. S1A–C). Furthermore, we averaged the coordinates of the lesion sites over the same window size.

To evaluate the motor deficit, we calculated the total number of arm entries and maximum moving speed between arms. The maximum moving speed was analyzed separately for the right and left selections at pretest and POD2. All behavioral analyses of video data were performed with a custom written Python program (https://www.python.org/) in Google Colaboratory (https://colab.research.google.com/notebooks/intro.ipynb). We used the results of the analysis conducted by two trained technical assistants (from our previous study [13]) for examining the inter-rater reliability of the analysis program (Supplemental Fig. S2 and Supplemental Table S1).

#### 2.3.2. Evaluation of motor paralysis

We used the ladder rung walking task for assessing motor paralysis after a focal cerebral infarction [20] (Fig. 4). The horizontal ladder apparatus and recording system were adapted from a previous study (Fig. 4B) [21]. Specifically, the horizontal ladder was 100 cm in length, 3.2 cm in width, and 56 cm in height from the ground. Two ladder conditions were used: regular pattern (alternating) and irregular pattern (random). In the regular pattern, the rungs were spaced 1 cm apart. In the irregular pattern, the spacing between rungs varied from 1 to 2 cm, with <3 rungs of 1 cm spacing. We generated 10 irregular-rung patterns using a spreadsheet (Microsoft Excel, Microsoft Corporation, USA), randomly selected and used in all trials. We recorded mouse movements from three directions (right, left, and down) using two mirrors and a digital video camera (C922; Logitech, Lausanne, Switzerland). All mice were transferred to the testing room to acclimate ≥1 h prior to each behavioral test. Training consisted of five trials/day on the ladder rung task (regular rung placement) over two consecutive days, starting three days before the focal photothrombotic infarction [22]. All mice were subjected to this pretest before photothrombosis. Two days thereafter, each mouse performed a total of six posttests. In the pre and posttests, each ladder condition (regular and irregular) was used in three trials.

To assess motor paralysis after a focal cerebral infarction, we calculated the error rate in the ladder rung walking task as missteps (error)/total steps (Fig. 4C). We defined a misstep as occurring when the tips of the mouse toe reached a position, which was 3 mm below the ladder height. All videos were recorded at 1920 × 1080 pixels and cropped to 300 × 900 pixels for further analysis. The video of the ladder rung walk task was analyzed by DeepLabCut, a markerless pose-detecting machine learning Python package [23]. We trained the neural network for 406,690 iterations on 172 labeled frames with the position of all four limbs, the nose, and tail for three side views. Limb missteps were detected by a custom Python program.

### 2.4. Nissl staining and slice analysis

To determine the infarction size, extent of lesions, and centroid of lesion areas (A-P direction), we stained brain slices (100 μm)/ with 0.1% cresyl violet acetate (Sigma-Aldrich Co., LLC. Tokyo, Japan) as previously reported [13]. Details were described in our previous study. We calculated all values using ImageJ software (https://imagej.nih.gov/ij/) and Matlab.

### 2.5. Statistical analysis

We used IBM SPSS, version 23.0 for windows and for all data analyses, and presented the data as mean ± standard error of the mean (SEM). Statistical significance was considered at *p* < 0.05.

## 3. Results

### 3.1. Photothrombotic infarction within AGm2.0, 1.5, 1.0, and 0.5

Fig. 1B illustrates examples of the infarct site in AGm2.0, 1.5, 1.0 and 0.5. The centroid of the lesion area (A–P direction) and lesion volumes on POD18 in all groups are shown in Fig. 1C. There were no differences in infarct volumes between groups (*F*(3, 52) = 0.243, *p* = 0.866; Fig. 1C).

### 3.2. Custom program reliability for the left-side selection rate

To assess ipsilesional spatial bias after focal cerebral infarction, the left-side selection rate determined by two trained technical assistants (Evaluators 1 and 2) and a custom Python program are presented in Supplemental Table S1. The inter-rater reliabilities for the left-side selection rate in the Control and Stroke groups among the two evaluators and the custom Python program were excellent, with intraclass correlation coefficients (ICC)_(2,1)_ ranging from 0.97 to 0.98 and from 0.98 to 0.99, respectively (Supplemental Table S1). Furthermore, there were positive correlations between Evaluator 1 and 2, between Evaluator 1 and program, and between Evaluator 2 and program in all data (Supplemental Fig. S2A–F). Thus, there were no inter-rate differences between the program and trained evaluators.

### 3.3. Behavioral changes after infarction

We collected the left-side selection rate, total number of arm entries and maximum moving speed at five time points: pre, acute (POD2 and POD3), mid-term (average of POD 4–7), and recovery period (average of POD 10–18) [13].

#### 3.3.1 Changes in left-side selection rate after infarction

There was a significant interaction between group and times for the left-side selection rate (*F*(13.379, 207.375) = 3.184, *p* < 0.001; Fig. 2B). In all groups except the control, the left-side selection rate at POD2 was significantly lower than that at pretest (*p* < 0.05; Fig. 2B). In the AGm1.0 and AGm0.5 groups, the left-side selection rate at POD10–18 was significantly increased (recovered) compared to that at POD2 (*p* < 0.05 and *p* < 0.01, respectively; Fig. 2B and C). In addition, in the AGm0.5 group, the observed decrease in the left-side selection rate returned to baseline levels by POD4–7 (*p* < 0.05). Conversely, in the AGm2.0 and AGm1.5 groups, there was no significant difference between the left-side selection rate at POD2 and POD10–18 (*p* > 0.05).

#### 3.3.2 Changes in total number of arm entries

We calculated the total number of times a given mouse entered an arm of the maze (Fig. 2D); no significant interaction was observed between group and time (*F* (11.887, 184.246) = 1.411, *p* > 0.05).

#### 3.3.3 Changes in maximum moving speed after infarction

We calculated the maximum moving speed after focal infarction (Fig. 2E); no significant interaction between group and time was observed (*F* (12.046, 186.719) = 0.835, *p* > 0.05). Furthermore, the maximum moving speed was analyzed separately for right and left-side selections at pretest and POD2. After cerebral infarction, the left-right ratio of the maximum moving speed for right and left-side selections was significantly lower only in the AGm1.0 group (*t* = 4.466, *df* = 13, *p* < 0.01; Fig. 2F and G).

### 3.4. Recovery rate of ipsilesional bias with respect to the infarction site and volume

We examined the correlation between the infarct site and degree of ipsilesional bias recovery, as well as between infarct volume and degree of ipsilesional bias recovery. In the acute phase, no correlations were observed between the infarct site and severity at POD2 (*r* = −0.107, *p* = 0.446; Fig. 3A). Conversely, the moving average of the infarction site and severity showed that the severity at POD2 was higher at 1 mm from the bregma (Fig. 3A). In the mid-term, we observed no correlations between the infarct site and recovery during the average of POD4–7 (*r* = −0.232; *p* = 0.095; Fig. 3B). In contrast, we found a significant negative correlation between the infarct site and recovery during the average of POD 10–18 (*r* = −0.294; *p* = 0.033; Fig. 3C). The moving average of infarction site and recovery showed that the recovery during the average of POD 10–18 was lower at 2 mm from the bregma (Fig. 3C). These findings, indicating that AGm lesions located more anteriorly resulted in less favorable recovery and support our previous findings [13].

**Figure 3.**
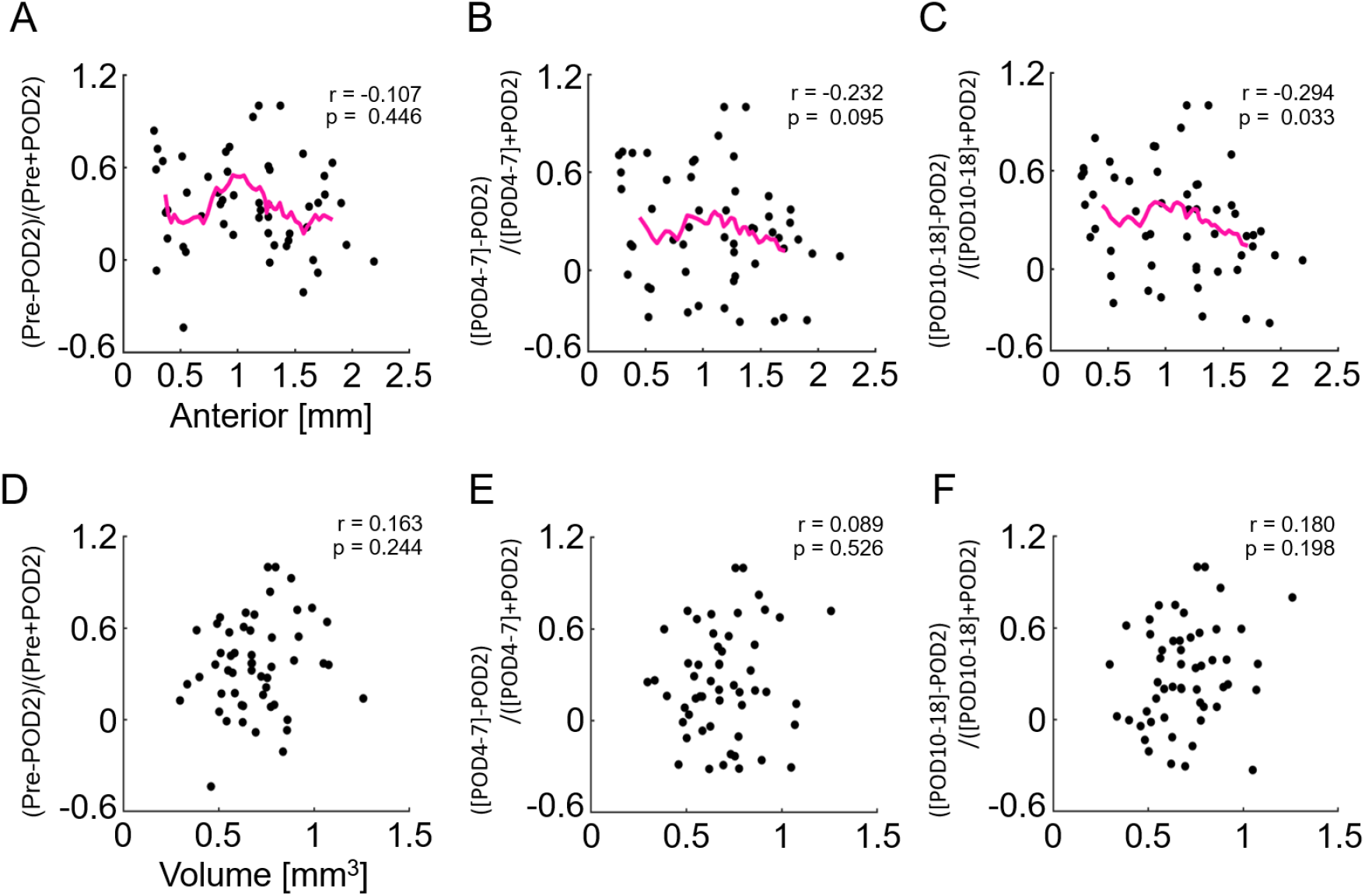
Relationship between infarction regions and changes in left-side selection rate, and between infarct volumes and changes in the left-side selection rate. A scatterplot of the infarct regions and severity; (pretest − POD2)/(pretest + POD2) at (A) the acute period. Scatterplots of the infarct regions and the degree of recovery; ([average POD4–7] − POD2)/([average POD4–7] + POD2) at (B) the mid-term and ([average POD10–18] − POD2)/([average POD10–18] + POD2) at (C) the recovery period. Infarct volume and severity at (D) the acute period. (E) Infarct volumes and degree of recovery at (E) the mid-term and (F) recovery periods. A high severity score in A and D indicates a high ipsilesional spatial bias. A large degree of recovery (B, C, E, and F) indicates a decrease in ipsilesional spatial bias. Correlations were identified using the Pearson’s correlation coefficient. *r* and *p* values are shown as indicated. The magenta line indicates the moving average.

In the infarct volume, there was no correlation between infarct volume and severity of ipsilesional spatial bias (*r* = 0.163, *p* = 0.244; Fig. 3D), between infarct volume and recovery at the average of POD 4–7 (*r* = 0.089, *p* = 0.526; Fig. 3E) or POD10–18 (*r* = 0.180; *p* = 0.198; Fig. 3F).

### 3.5. AGm infarction did not induce motor paralysis

To assess motor paralysis after focal infarction, the error rate in the ladder rung walk task was calculated before and after cerebral infarction. The centroid of the lesion area (A–P direction) and volumes at POD7 in all groups are shown in Fig. 4A. There was no difference in infarct volume among groups (*F*(3, 41) = 0.629, *p* = 0.601; Fig. 4A). For the hindleg error rate in the irregular condition in the AGm 2.0 group, there was a significant interaction between the left-right difference in the percentage of limb off the ladder (error rate) and time (*F* (6, 84) = 3.073, *p* < 0.01; Fig. 5). The error rate of the right hindpaw was higher than that of the left hindpaw at pretest (*p* < 0.05). Conversely, there was no interaction between left-right differences in error rate and time in other groups or conditions (*p* > 0.05).

**Figure 4.**
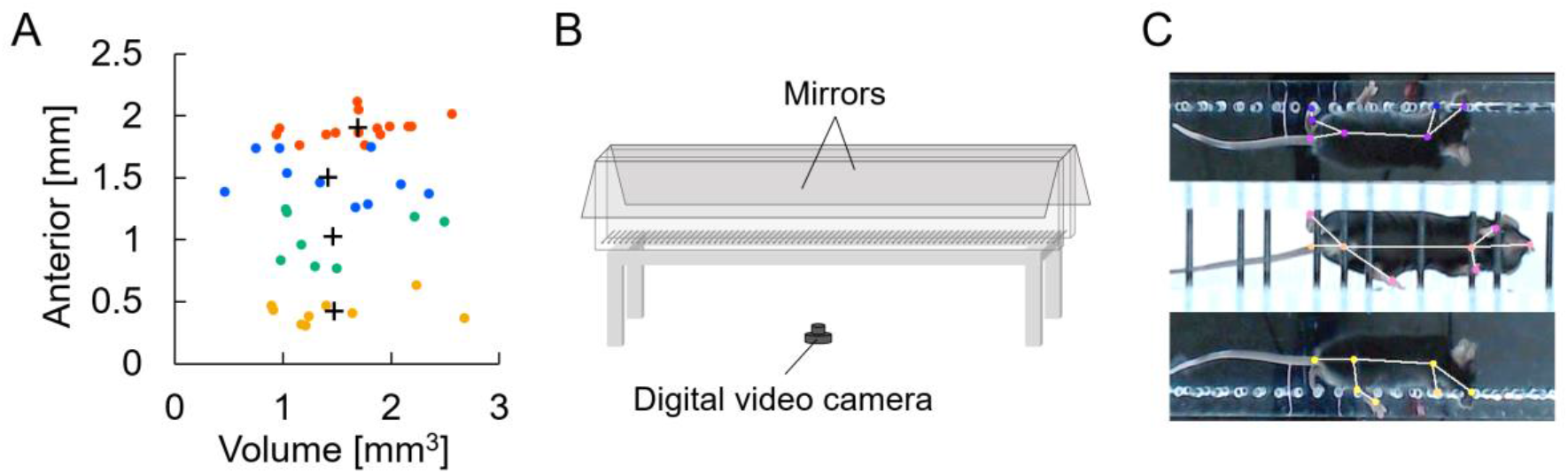
Photothrombotic infarction and experimental set-up of the ladder rung walking task. (A) A scatterplot of the infarct site and volumes at postoperative day (POD) 18. + indicates the average centroid of the lesion area (A–P direction) in the AGm2.0 (n = 15), AGm1.5 (n = 10), AGm1.0 (n = 8) and AGm0.5 groups (n = 9) and the average infarct volume in each group. The horizontal axis indicates the distance from midline to lateral and the vertical axis indicates the distance from bregma to the anterior end. Correlations were identified using the Pearson’s correlation coefficient. For a nonlongitudinal analysis of the infarction volume we used a one-way ANOVA. (B) Schematic diagram of the experimental apparatus. (C) Example of stepping off a ladder and markers of the area chosen for postural analysis.

**Figure 5.**
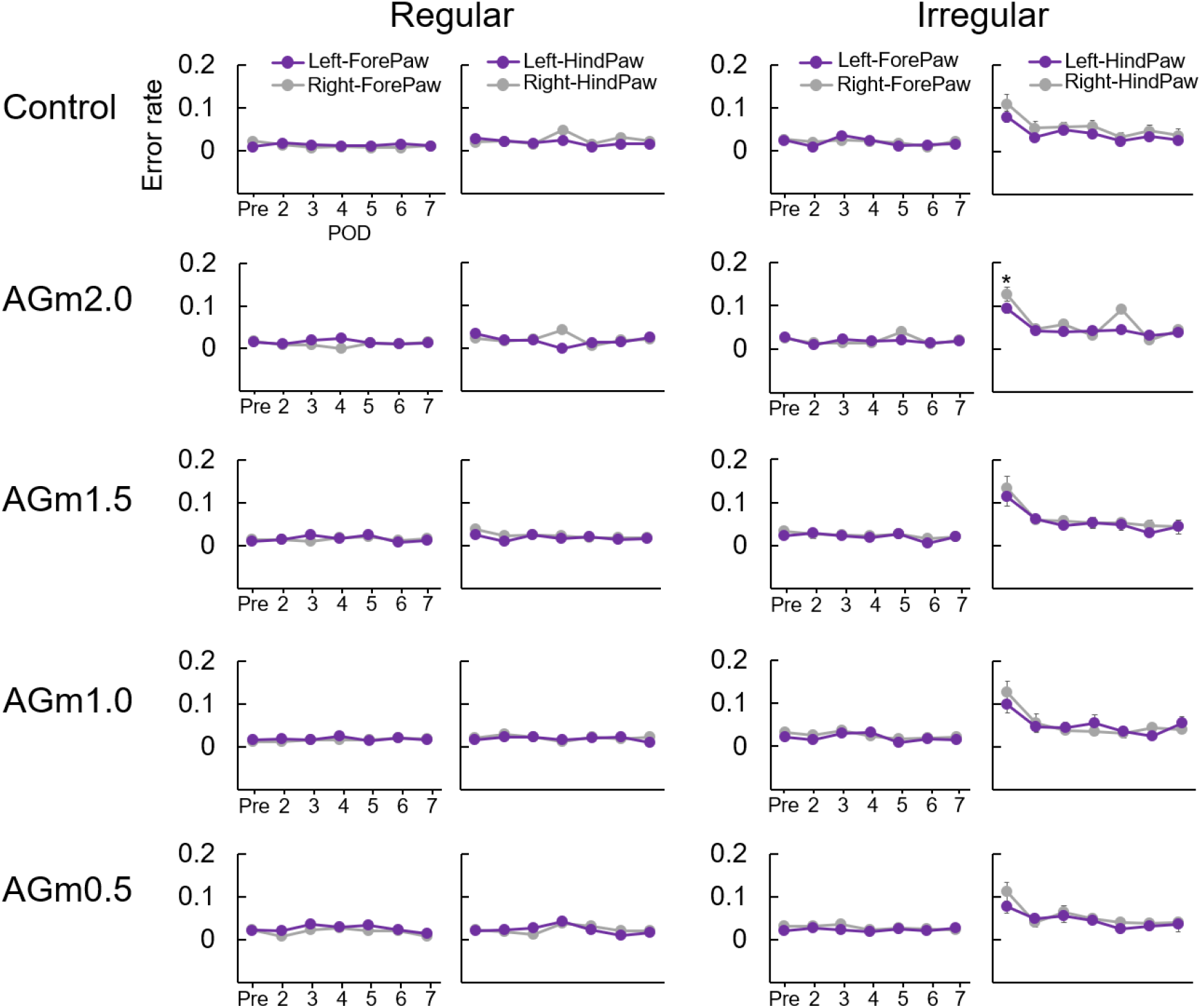
Evaluation of motor paralysis using the ladder rung walking task. Time course of error rate for the left and right forepaw and left and right hindpaw in the ladder rung walking task under regular and irregular conditions. *p < 0.05, Left paw vs. Right paw at each time point by two-way repeated measures ANOVA followed by Bonferroni’s test.

## 4. Discussion

This study yielded the following key findings: (1) inducing a unilateral local photothrombotic infarction in AGm2.0, AGm1.5, AGm1.0, or AGm0.5 (2.0 mm, 1.5 mm, 1.0 mm, and 0.5 mm anterior to the bregma, respectively) resulted in USN-like behavior in the acute phase, depending on somatosensory information; (2) the severity in acute phase was higher when the lesion was located 1.0 mm from the bregma; (3) AGm lesions located more anteriorly resulted in less favorable recovery; (4) no motor paralysis was observed in any of the mice with lesions in each AGm; and (5) the developed USN-like behavior analysis program showed high inter-rater reliability. These results indicate that there are functional differences within the AGm regions in spatial attention.

The AGm plays an important role in integrating sensory information with motor actions and decision-making processes [10–12]. Previous studies have shown differences in sensory modality-specific innervation, such as visual and somatosensory, from rostral to caudal within AGm [15, 16]. Therefore, here, we investigated the functional differences between different AGm regions in terms of acute symptoms and recovery of USN-like behavior. The results showed that all unilateral focal infarctions in the AGm (AGm2.0, AGm1.5, AGm1.0, or AGm0.5) decreased the left selection rate. However, there was no significant difference in the error rate between the left and right paw in the ladder rung walking task after unilateral focal infarction in the AGm. These results indicate that all unilateral focal infarctions in the AGm induced USN-like behavior without motor paralysis. Among the various AGm regions, acute severity was higher when the lesion was positioned 1.0 mm from the bregma (Fig. 3A). Additionally, after cerebral infarction, the left-right ratio of the maximum moving speed for right and left-side selections was significantly lower only in the AGm1.0 group (Fig. 2F and G). These results suggest that the intermediate caudal–rostral position of AGm may play a significant role in direction selection.

A previous study suggested that the rostral and caudal AGm play different roles in directed attention, as the neglect resulting from lesions in the caudal AGm is more profound and longer lasting than that from more rostral lesions [14]. This result is inconsistent with the present results. One possible reason for this difference is that there are differences in the sensory modality of neglect evaluated in each study. The previous study used a task that depended on visual information to evaluate USN-like behavior. In contrast, in our study, we used a task that relied solely on somatosensory information without visual cues and found delayed recovery from USN-like behavior in rostral AGm lesions, as this area has numerous connections to the somatosensory cortex [15, 16]. These findings suggest that the recovery from USN-like behavior may be influenced by the specific neural control features of the AGm.

We examined the role of the rostral of AGm (AGm2.0) in the recovery of USN-like behavior after local photothrombotic infarction. The findings from this study, indicating that AGm lesions located more anteriorly resulted in less favorable recovery and support our previous findings [13]. Interestingly, previous research indicated the potential for increased recovery by manipulating specific cellular processes within the rostral AGm. For instance, overexpression of cAMP response element binding protein (CREB) in a subset of peri-ischemic neurons in this area enhanced axonal sprouting and improved behavioral recovery after stroke [24]. Another recent study has reported that inhibition of phosphodiesterase 2A, which attenuates CREB activation, in the rostral of AGm can enhance functional recovery, increase axonal projections in the peri-infarct cortex, and enhance the functional connectivity of motor system excitatory neurons [25]. Together with our results, CREB activation in the rostral AGm seems to plays a crucial role in recovering from neglect after damage. Future research should focus on identifying the specific cellular and molecular mechanisms promoting recovery and exploring potential interventions targeted at enhancing plasticity within this area to improve recovery outcomes.

The current study has a limitation. We investigated the relationship between the location of AGm lesions and severity of symptoms depending on somatosensory information. The recovery from USN-like behavior tended to be slower in cases with more rostral lesions in the AGm, suggesting a potential for long-term persistence (severity) of neglect in specific sensory modalities associated with plastic compensation of the connected areas. However, as we did not evaluate UNS-like behavior related to visual information, the relationship between the location of AGm lesions and severity of symptoms depending on visual information remains unclear.

In summary, this study aimed to investigate the functional differences across AGm subregions in terms of acute symptoms and recovery characteristics of USN-like behavior in mice. In the acute phase, the severity was higher when the intermediate caudal–rostral position of the AGm was lesioned. This result suggests that the intermediate caudal–rostral position of AGm may play a significant role in direction selection. In contrast, the recovery from USN-like behavior tended to be slower with rostral lesions, suggesting that the functional recovery depends on the connected areas.

## Author contributions

DI, HO, AY, and TY conceived and designed the experiments. DI, HO, and TY collected and analyzed the data. DI, HO, TY, SY, and MM wrote the manuscript. MM and YK supervised all studies. All authors reviewed and revised the manuscript.

## Declaration

The authors declare no competing interests. All experiments were performed in accordance with the “Guidelines for the Proper Conduct of Animal Experiments” established by the Science Council of Japan in 2006, and were approved by the Animal Care and Use Committee of Ibaraki Prefectural University of Health Sciences (approval number 2022-14).

