## Supplementary information for "Role of the medial agranular cortex in unilateral spatial neglect"

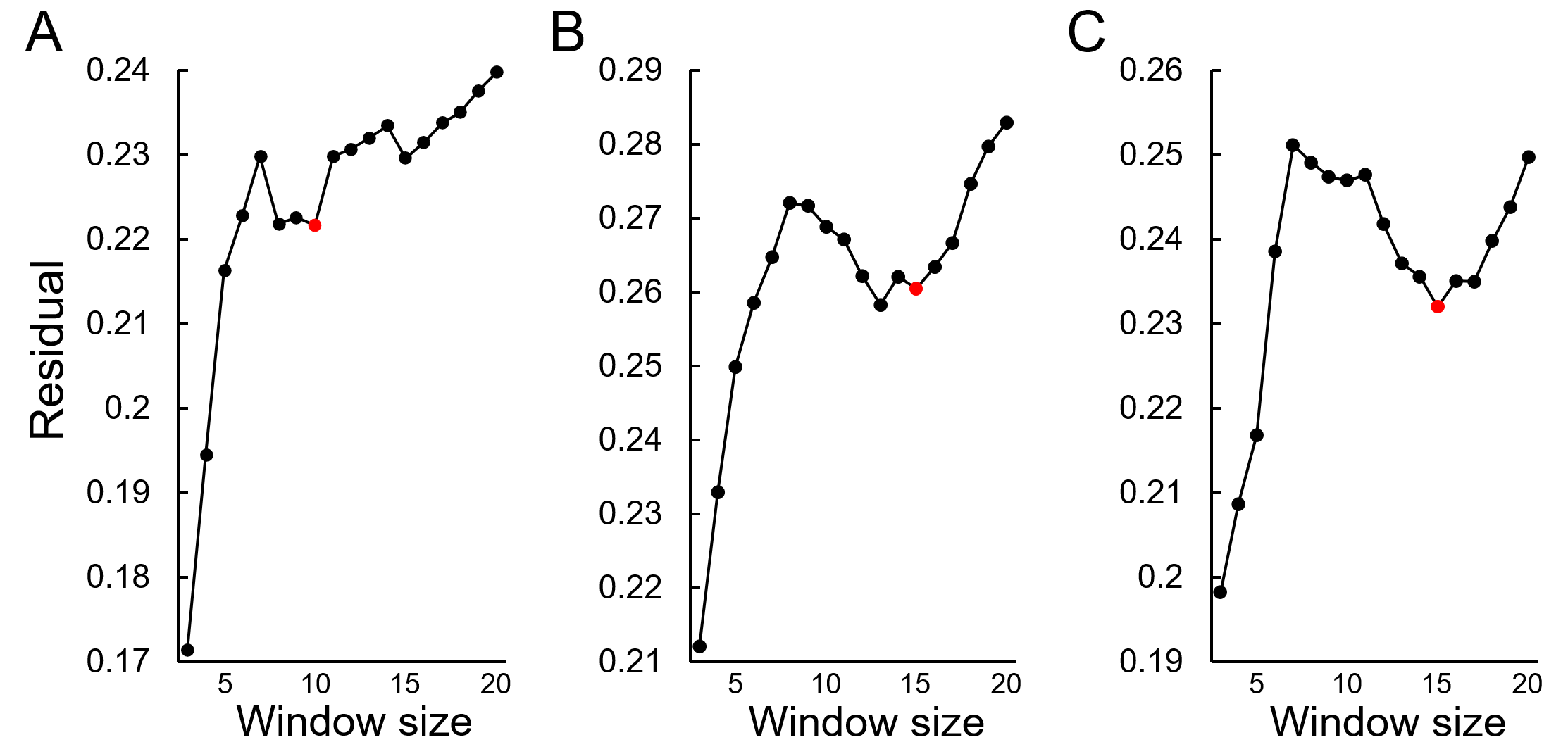

**Supplementary Figure S1. Relationship between sample window size and residuals for moving averages related to Figure 3**

(A) The severity at POD2. The red dot indicates the window size used in Fig. 3A. (B) The degree of recovery at mid-term (POD4-7). The red dot indicates the window size used in Fig. 3B. (C) The degree of recovery at recovery period (POD10-18). The red dot indicates the window size used in Fig. 3C.

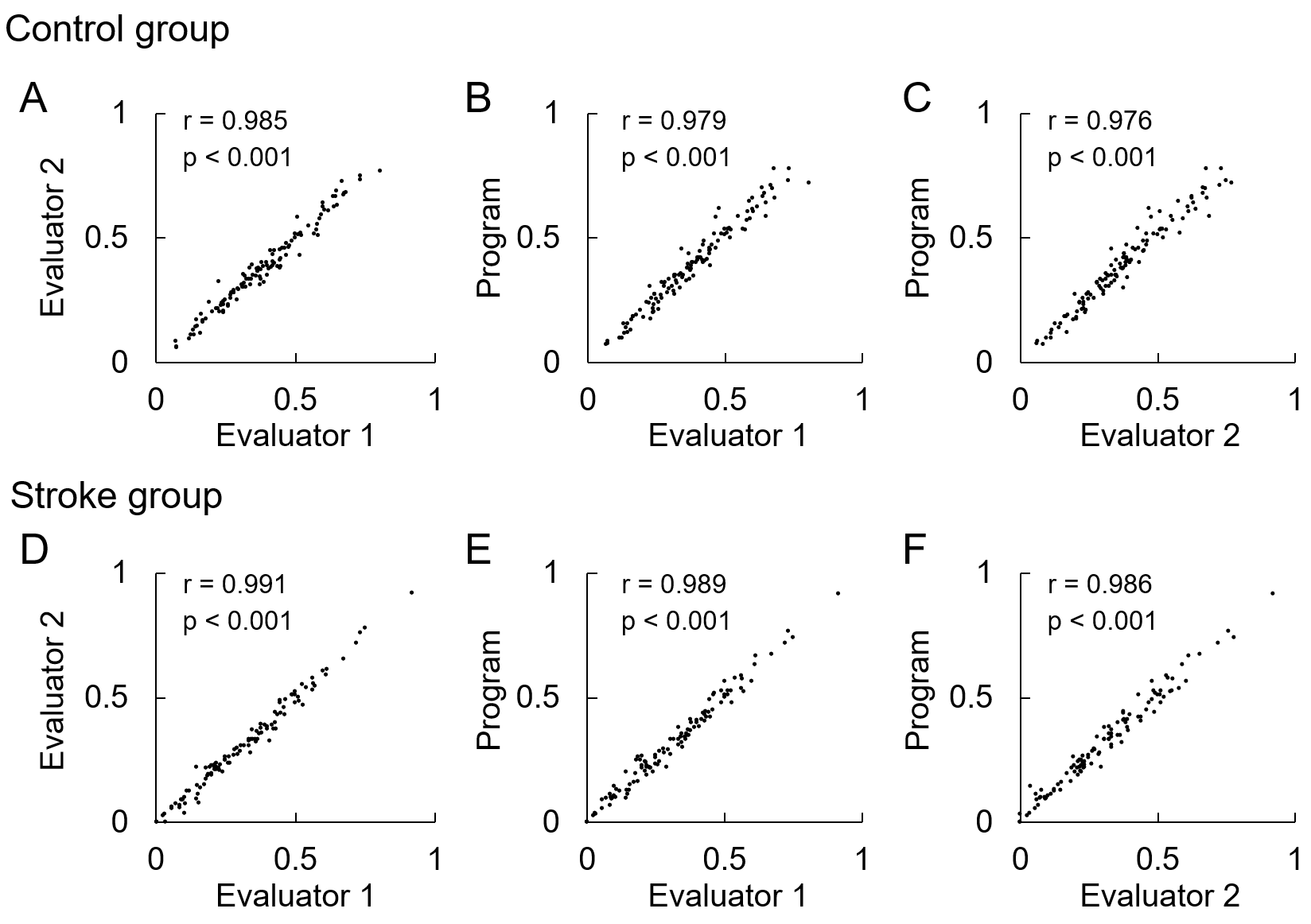

**Supplementary Figure S2. Inter-rater reliability for left-side selection rate**

Inter-rater reliabilities for the left-side selection rate in (A–C) Control and (D–F) Stroke groups among the two evaluators and the custom Python program. Correlations were identified using the Pearson’s correlation coefficient.

|  |  | Left-side selection rate Men (SEM) | | | | | | | | | | | | | | | | | | | |  | ICC (95% CI) |  | *P* |
| --- | --- | --- | --- | --- | --- | --- | --- | --- | --- | --- | --- | --- | --- | --- | --- | --- | --- | --- | --- | --- | --- | --- | --- | --- | --- |
|  |  | Pre | | POD2 | | POD3 | | POD4 | | POD5 | | POD6 | | POD7 | | POD10 | | POD14 | | POD18 | |  |  |  |  |
| Control | |  |  |  |  |  |  |  |  |  |  |  |  |  |  |  |  |  |  |  |  |  |  |  |  |
|  | Evaluator 1 | 0.34 | (0.05) | 0.38 | (0.05) | 0.40 | (0.05) | 0.40 | (0.06) | 0.35 | (0.06) | 0.44 | (0.04) | 0.40 | (0.06) | 0.37 | (0.04) | 0.40 | (0.03) | 0.35 | (0.03) |  | 0.98  (0.97, 0.98) |  | <0.001 |
|  | Evaluator 2 | 0.33 | (0.05) | 0.37 | (0.05) | 0.40 | (0.05) | 0.40 | (0.06) | 0.36 | (0.05) | 0.43 | (0.05) | 0.40 | (0.06) | 0.36 | (0.05) | 0.39 | (0.03) | 0.35 | (0.03) |  |  |  |  |
|  | Program | 0.35 | (0.06) | 0.37 | (0.06) | 0.41 | (0.05) | 0.40 | (0.05) | 0.37 | (0.06) | 0.45 | (0.05) | 0.42 | (0.06) | 0.39 | (0.05) | 0.41 | (0.03) | 0.35 | (0.03) |  |  |  |  |
| Stroke | |  |  |  |  |  |  |  |  |  |  |  |  |  |  |  |  |  |  |  |  |  |  |  |  |
|  | Evaluator 1 | 0.43 | (0.04) | 0.16 | (0.04) | 0.28 | (0.05) | 0.33 | (0.06) | 0.38 | (0.06) | 0.37 | (0.06) | 0.25 | (0.05) | 0.35 | (0.06) | 0.33 | (0.06) | 0.32 | (0.06) |  | 0.99  (0.98, 0.99) |  | <0.001 |
|  | Evaluator 2 | 0.42 | (0.04) | 0.17 | (0.04) | 0.27 | (0.05) | 0.31 | (0.06) | 0.37 | (0.07) | 0.36 | (0.06) | 0.24 | (0.05) | 0.35 | (0.06) | 0.32 | (0.05) | 0.32 | (0.06) |  |  |  |  |
|  | Program | 0.43 | (0.04) | 0.18 | (0.04) | 0.28 | (0.05) | 0.33 | (0.06) | 0.37 | (0.07) | 0.37 | (0.06) | 0.26 | (0.05) | 0.36 | (0.06) | 0.33 | (0.05) | 0.33 | (0.06) |  |  |  |  |

**Supplementary Table S1.** Descriptive statistics and intraclass correlation coefficients (ICC)_(2,1)_ of the analysis results by Evaluator 1, 2, and custom Program
